# Quantitative model and physical mechanisms of iRBC membrane curling during egress of malaria parasites

**DOI:** 10.1101/2024.02.13.579773

**Authors:** N. Gorkavyi, A. Parmeggiani, A. V. Kajava

**Affiliations:** Science Systems and Applications Inc., Lanham, MD; Laboratory Charles Coulomb, University of Montpellier, CNRS UMR 5221, Montpellier, France; Centre de Recherche en Biologie cellulaire de Montpellier, University of Montpellier, CNRS UMR 5237, Montpellier, France

**Keywords:** Malaria, merozoite egress, physical mechanism, Plasmodium genus, quantitative model, red blood cell

## Abstract

Egress of malaria merozoites from infected red blood cells (iRBC) is a critical step in the parasite’s life cycle. The egress is accompanied by the formation of a pore in the erythrocyte membrane, followed by an outward curling of the membrane around the pore, resulting in a complete eversion of the erythrocyte membrane, pushing the parasites away. Despite the well-documented steps of the egress, the detailed mechanism and source of energy for such a spectacular eversion of iRBC remains largely unknown. In this paper, we consider a biophysical model based on the energetics of the egress process that includes both viscous dissipation and energy consumption for the formation of the rim around the pore in iRBC. We show that viscosity does not play a significant role in iRBC eversion and we hypothesize that this process is controlled by lateral lipid diffusion. The model is supported by quantitative estimates and is in good agreement with known experimental data.

## 1. Introduction

Malaria is one of the most important and widely distributed tropical diseases, which kills hundreds of thousands of people yearly (in 2017, ∼435,000 people died of malaria [1]). The malaria parasite of the *Plasmodium* genus replicates within erythrocytes, producing merozoites that are released from infected red blood cells (iRBC) via a poorly understood process called egress [2-7 and references therein]. The cause of the iRBC rupture and egress has long been considered as a result of an overpressure produced by merozoites inside the cell, however, recently, Braun-Breton and colleagues have shown that the mechanism of the rupture is much more complicated and consists of several steps [8-10]. First, a small pore is formed in the iRBC membrane, through which, due to overpressure, 1-2 merozoites come out. This opening equalizes the pressure between the cell interior and the external environment. Other merozoites remain in the cell. Second, the cell membrane begins to turn inside out, forming a rim around the pore (Figure 1) and expanding the initial pore to the diameter of the red blood cell (Figure 1c). Third, the membrane continues to curl until a complete eversion and ejection of all merozoites into the extracellular space. As a result, the cell is transformed from a sphere with a pore to an open disk with a toroidal rim (Figure 1d). Interestingly, from geometrical considerations, one can argue that the elastic energy stored in the membrane before and after the egress increases (Figure 1). The elastic energy stored in a bent membrane is calculated using a cell membrane typical bending modulus k, and the radius of curvature of the bent membrane. For example, a membrane rolled into a ball (see Fig. 1a) has an elastic energy 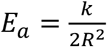. Let us compare the shapes of the membrane in Fig. 1a and 1d. The smaller R, the greater the elastic energy stored in the membrane. Obviously, the membrane in Fig. 1d has a much smaller radius of curvature than in Fig. 1a. If we assume the same elastic properties of the membranes in both states, then it can be shown (see Appendix II) that the elastic energy of the membrane in Fig. 1d is significantly greater than in Fig. 1a: *E*_*a*_< *E*_*d*_. We come to the paradoxical conclusion that the elastic energy of the erythrocyte in the process of eversion increases, which, at first glance, makes this process energetically unfavorable and forbidden. Hence, the assumption that membrane properties are the same in both states in Figs. 1a and 1d is incorrect. Obviously, the iRBC membrane in Fig. 1a has a latent source of energy *E*, which is spent both on the tight twisting of the membrane and on *E*_*visc*_, i.e., the energy losses to overcome viscosity when the membrane moves in a liquid medium. The energy balance can be written as:

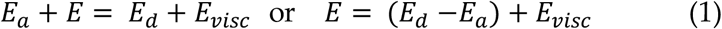

Below we will discuss the contribution for *E* and possible sources of energy in the membrane of an infected erythrocyte: spontaneous tension, cytoskeleton, attachment of small vesicles, etc. It is obvious that a quantitative analysis of these latent energy sources is difficult, because even their nature is unclear. However, we can use the energy equation (1), which equates energy sources and sinks. Energy sinks are easier to calculate: they include both an increase in the elastic energy of the coiled membrane (*E*_*d*_−*E*_*a*_) and various dissipative contributions, for example due to blood viscosity *E*_*visc*_.

**Figure 1.**
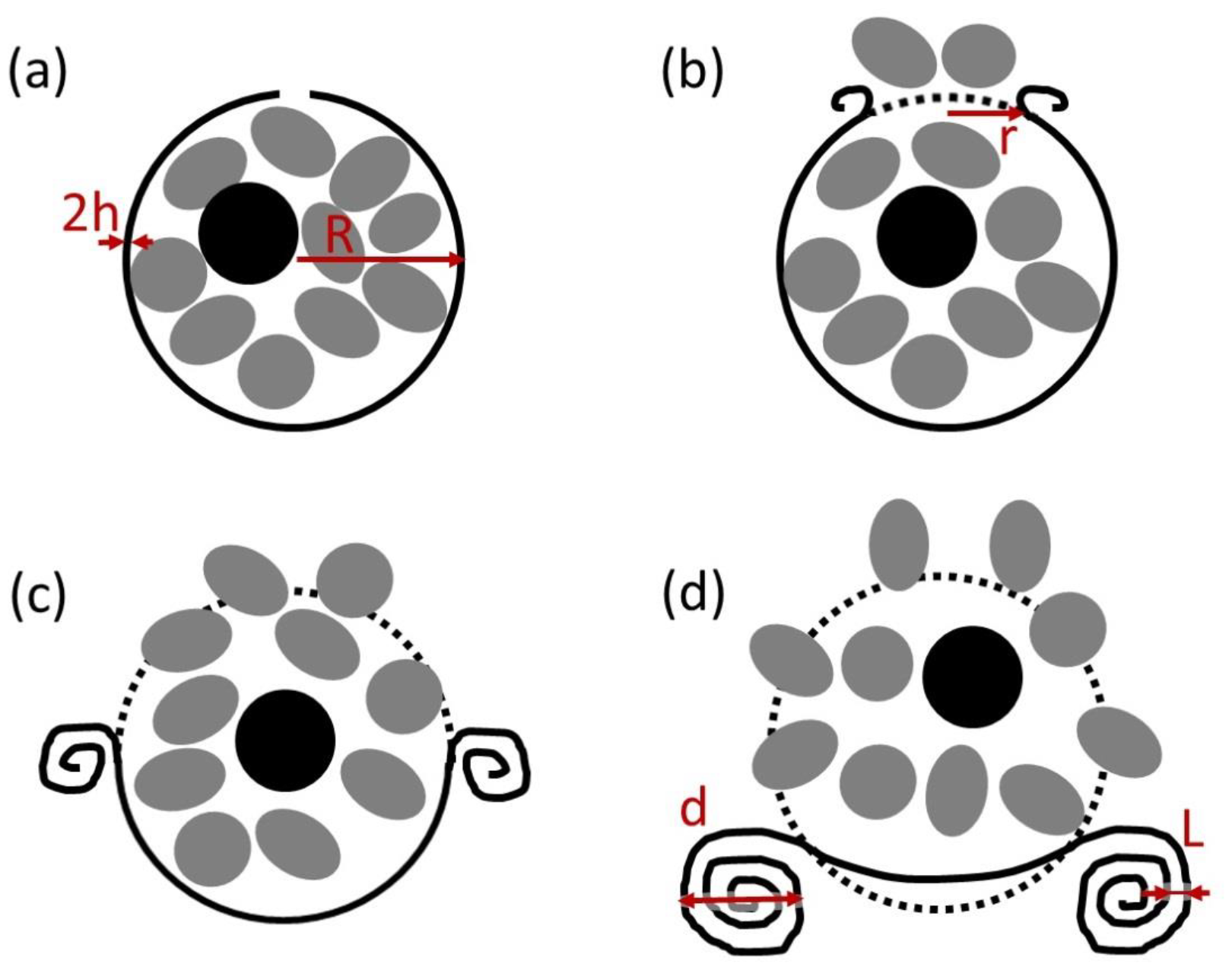
A schematic representation of (a) the iRBC after a pore opening; (b) partially opened erythrocyte; (c) half-opened cell and (d) a complete eversion of the iRBC. Merozoites are marked with gray ovals; black spot corresponds to the food vacuole [1,8]. Red arrows point to the rim of the folded membrane. Parameters used to describe the process of membrane deformation are: ***h*** *is the* half-thickness of the membrane, ***R*** is the radius of the cell, ***r*** is the radius of the pore, ***d*** is the diameter of the torus cross-section, and ***L*** is the distance between adjacent membranes in the roller.

The erythrocyte membrane is two-layered; therefore, for the spherical shape in Fig. 1a (see also Appendix I) the area of the inner layer should be less than the area of the outer layer. However, through the process of eversion of the erythrocyte, the inner layer of the membrane becomes the outer one. In order for this process to take place without ruptures of the inner layer, it is necessary that its area increases by 2% (see Appendix I). Hence, we can assume that the inner layer of the iRBC membrane in the initial state (Fig. 1a) has an excess of energy and matter (for example, lipids), condition which ensures the process of eversion of the erythrocyte shown in Fig. 1.

What does determine the characteristic restructuring time of the iRBC membrane? Obviously, merozoites inhibit the release of energy *E* for a considerable time, triggering the process of cell eversion only at the right moment. We can hypothesize that after the start of the process, its rate is no longer controlled by the influence of merozoites, but by other factors such as, and most likely, the lateral rate of lipid diffusion. The experimentally measured iRBC opening time [8] coincides with the lipid diffusion time at a distance of the order of the cell radius *R*. We could slower or stop the movement of lipids, for example, by cooling the system. Then the restructuring of the cell (see Appendix I and II), including the elastic expansion of the pore, would be difficult or even impossible to accomplish, making the process of egress inefficient or even stopping it. In these conditions, even though the energy *E* stored in the inner layer of the membrane would still exist, but its consumption would be inhibited due to the low mobility of lipids. It is possible that other factors control the process of iRBC opening such as, for example, the characteristic time of cytoskeleton rearrangements on the inner layer of the membrane, but the evaluation of the role of other factors requires additional research that is beyond the scope of this article.

A question arises: What is the source of energy for the iRBC transformation? In a previously published model, Abkarian et al. (2011) associate cell eversion with an excess of elastic energy in the overstressed internal leaflet [8]. The source of this energy excess however remains unclear. This model is based on the equality between elastic energy consumption and viscous dissipation. In this case, the curling time of a cell should be proportional to viscosity. As noted by Abkarian et al. [8], the observed curling time is two orders of magnitude smaller than the value based on viscous dissipation obtained from theoretical calculations. The agreement between the experiment and the theory of Abkarian et al. [8] requires that the viscosity used in the model be about a hundred times greater than the normal blood viscosity. This indicates the incompleteness of the model and the need to take into account new effects.

In this paper, we first propose a quantitative model that includes not only viscous dissipation but also energy consumption that drives the process of formation of the rim around the pore in iRBC.

Unlike previous models, we are discussing here the origin of the possible sources of energy and matter related to the shape change of an iRBC prior to and after merozoites egress. Eversion of the cell and expansion of the pore from size *r* to *R* cannot occur without a strong stretching of the membrane with a dilation ratio coefficient *R*⁄*r*. Obviously, either energy-consuming membrane ruptures or a more global and less energetically expending rearrangement of the lipid layer accompany the membrane stretching. In this transformation, lipids of both layers move across the membrane so that it can stretch and produce an eversion without tearing. The role of lateral lipid diffusion as the main factor limiting the rate of eversion of infected RBC has actually not yet been studied. In doing so, we also provide a reasoned estimate of the amount of matter necessary to prepare the iRBC mechanical instability prior the pore formation. These results are the main new outcome of this work, thus providing a new physical explanation that can be matter of future experimental tests of an important process such as the release of malaria parasites from an infected cell.

The manuscript is organized as following. In section 2, we formulate the model employed, discussing the equation that provide the total energy balance of the cell membrane and from which we obtain the equation of the time evolution of the membrane during its eversion. From this equation, we show and discuss the main contributions of energy loss and transduction during the membrane dynamics in agreement with experimental estimates and measurements. Section 3 is devoted to the results obtained and the discussion of the possible origins of the energy and matter excess that drive the membrane eversion. We then discuss iRBC eversion from possible scenarios as well as experimental evidences in agreement with our model results. Section 4 concludes the article. Several appendices detail the estimates employed in the main text.

## 2. Methods

### 2.1. Model formulation

In order to produce the observed process of eversion of an infected blood cell, it is necessary that an amount of energy *E* has been accumulated inside the cell prior its large shape transformation. The origin of this energy *E* is still poorly understood, although some of this energy may be stored in spontaneous curvature (we will discuss possible sources of *E* in Section 3). Even without knowing the precise origin of the sources of energy necessary for cell eversion, one can investigate the question: What is the stored energy E used for? In this article, we will focus on analysing the different contributions to energy consumption that are associated with the observable transformation of the cell and the movement of the folded membrane.

### 2.2. Energy consumption contributions

All the energy consumption contributions of the iRBC during the transformation can be divided into two classes: viscous and non-viscous. Viscous dissipation of energy during the motion of the rim in the viscous medium (with Stokes friction *F*_*Stokes*_) [11] writes as:

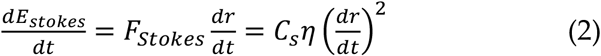

where *r* is the major radius of the torus (or the pore) evolving in time and *η* is the viscosity of the medium. For a cylinder with diameter *d* and length 2*πr* (see references in [12):

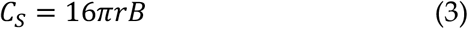

where 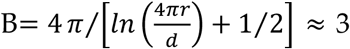 for *r* = 1 − 3*µm* and *d* = 0.4 − 0.8*µm*. We will use such an approximation in the following.

During eversion, the iRBC first consumes the stored energy in the formation of a rim and in viscous losses during the rim expansion. Non-viscous losses are then associated with an increase in the elastic energy of the folding membrane as well as the energy consumption for producing eventually the plastic deformation and rupture of the membrane. The elastic energy of the tightly folded multilayer membrane can be estimated (see Appendix II) as

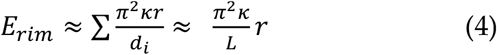

where κ is a cell membrane typical bending modulus; *d*_*i*_ – the diameters of the cylinders in the rim; *L* ≈ 0.1*µm* is the average distance between the layers of the rim (see Figure 1).

### 2.3. Energy balance

To obtain an equation describing the dynamics of erythrocyte eversion, we differentiate the energy equation (1) with respect to time. The total energy consumption 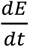 during the cell transformation, including both viscosity and elastic contributions, can be described by the sum of viscous dissipation 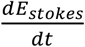 from Eq. (2), while non-viscous losses 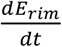 from Eq. (4), where we consider a cylindrical shape at the place of toroidal one (i.e. we neglect the main curvature radius of the toroid), by:

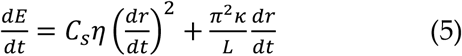

where *η* ≈ 10^−3^ *kg*⁄(*m* ∗ *s*) is the viscosity of the external watery environment and *C*_*s*_ ≈ 48*πr*. Assuming *r* ∼1 *µm*, it is possible to plot the dependence of the terms in equation (5) on the rate of erythrocyte unfolding (Fig. 2).

**Figure 2.**
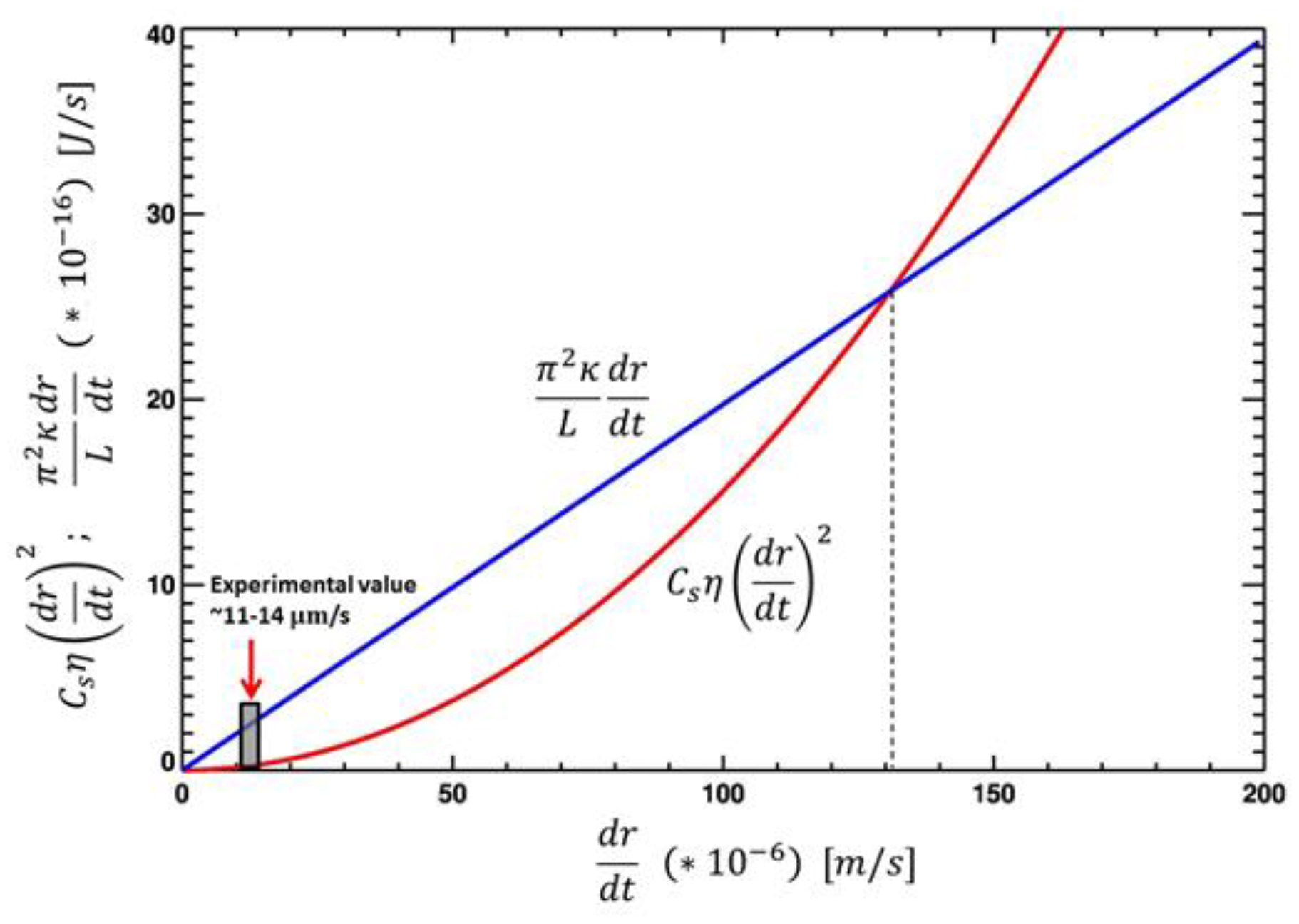
Dependence of the terms of equation (5), respectively the energy consumption for the elastic deformation (in blue) and the viscous dissipation (in red), on the rate of a hole opening in iRBC. The influence of viscosity and elasticity is equal at velocity 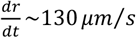. At lower speeds, the energy consumption for elastic deformation (in blue) dominates; at high speeds, viscous dissipation (in red) predominates.

The experimentally observed speed of the cell opening is 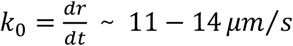, and the characteristic time for the curling stage is *t*_*curl*_∼ 0.1 − 0.2 *s* [8]. For such slow speeds 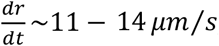, the energy consumption for elastic deformation will significantly exceed the energy consumption for viscous deceleration (see the following Section 2.4 Estimations).

### 2.4. Estimations

The model by Abkarian et al. (2011) [8] corresponds to the case when the right-hand side of Eq. (5) contains only the first viscosity term. In our model the influence of viscosity and elasticity is equal at velocities 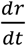 10-fold higher - see Eq. (5) and Fig. 2 - than those observed in experiments:

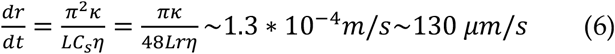

For speed 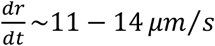 and *r* ∼1 *µm* we can estimate from Eq. (5) the intensity of the term with viscosity as 2 ∗ 10^−17^ *J*⁄*s*.

Let us estimate the relative value of the second term on the right-hand side of Eq. (5), dividing it by the first term with viscosity. For the second term with the rim elasticity, we get

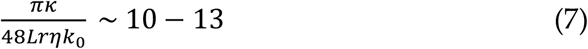

where *r* ∼1 *µm* and κ ≈ 2 ∗ 10^−19^ *J* [8].

Thus, the viscosity term is significantly smaller in magnitude than the term describing the time evolution of the rim radius on the right-hand side of Eq. (5). Note that in a number of papers noticeably lower speeds K_0_ of opening of the iRBC are observed [5-7]. With a decrease in the speed K_0_, the estimate Eq. (7) will increase even more, because the viscosity has a smaller effect on the slower movements of the membrane.

## 3. Results and Discussion

### 3.1. The contribution of viscosity and elasticity

From these estimates (see Eqs. 7-8), we conclude that viscosity does not play a significant role in the iRBC eversion [13]. The elastic deformation of the membrane and its rupture are more important and dominate the energy conversion/consumption process. The main contribution to energy consumption may not be the bending deformation of the membrane, but its ruptures. The relative role of a membrane rupture (or its stretching/compression) and its elastic deformation during bending can be clarified, for example, by experiments with a change in a cell membrane bending modulus κ.

*3.2. Role of lateral lipid diffusion*

We previously used the observed cell eversion rate 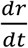 to compare the right terms of equation (5). Equation (5), however, does not allow a theoretical determination of 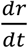, because the rate of energy consumption 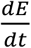 remains unknown. The greater 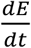, the greater the speed 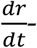 and vice versa. What does determine the rate of the excess energy consumption 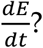 In order to explain the role and contribution of lipid rearrangement in the membrane leaflets, let’s carry out the following thought experiment: one can inhibit the lateral diffusion of lipids. Immobile lipids have no effect on energy consumption; they completely prevent cells from opening, except in the case of membrane rupture, but with much greater energy to tear the membrane leaflets. The example of a paper sphere is a good analogy for a cell with immobile lipids. It is obvious that such a ball cannot be turned out without numerous breaks. However, in case that at least a slow diffusion of lipids is allowed, then eversion of the cell without ruptures will be possible, but the rate of this eversion will be determined by the speed of lipid movement in the regions with maximal stress or potential ruptures. The faster the lipids move, the faster the cell turns inside out. The motion of lipids is governed by Brownian motion. The possible energy consumption *E* for the movement of lipids to the zones of maximal stress will be relatively small, because it belongs to the category of viscous losses, which are small at the observed opening rates.

We hypothesize here that the time of curling of iRBCs depends essentially on the rate of lateral diffusion of lipids in the lipid layer of the membrane. Indeed, an estimate of the typical time for a lipid to diffuse in the membrane on a typical distance *l*_*d*_∼ 1 *µm* gives:

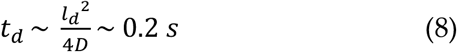

where *D* ∼10^−12^ *m*^2^⁄*s* is a typical value of the lateral diffusion coefficient of lipids (see, for example, [14-17]). The value *l*_*d*_ ∼ 1 *µm* characterizes the typical length of the diffusion or/and drift path of the lipid that travels to ensure the deformation of blood cells with a radius of 3 μm. Various factors can significantly slow down the diffusion of lipids, which will lead to the inhibition of the cell eversion process. This hypothesis can be tested experimentally by examining cell eversion at different lipid diffusion coefficients.

### 3.3. Possible Energy Sources

Our estimation yields a value of the energy consumption equal to 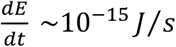 and total energy of 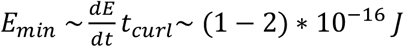 required for eversion of one cell. This value is minimal and can be noticeably higher if a significant part of the energy is used for the rupture of the membrane and the membrane-associated-cytoskeleton. For the following estimations, we assume that the transformation of the cell may require a maximal energy *E*_*max*_∼ 10^−15^ *J*. A question arises: What is the source of such required energy?

There is a set of mechanisms for changing the curvature of the membrane and its elastic energy, up to its rupture and vesiculation [3, 18-22]. These mechanisms are associated with the rearrangement of the cell cytoskeleton [23], with changes in the chemical composition of the membrane due to the attachment of lipids and proteins [22,24], as well as due to the attachment of macromolecular clusters such as vesicles to the membrane [25].

Let us consider in more detail the organization of the iRBC. The erythrocyte membrane consists of two lipid layers (inner and outer). We assume that the outside area of the membrane is not changed during cell membrane dynamics. The lipid exchange between the inner and outer layers is very slow (more than six hours, see, for example, a review by [26]) compared to the membrane deformation and therefore this event can be neglected during the fast curling phase. The elastic energy stored in the tightly rolled rim of the multilayer membrane is considerably larger than the initial elastic energy contained in the membrane of the spherical cell (see Appendix II). The energy cost for the viscous friction at the cell opening worsens the problem of the energy shortage.

### 3.4. Possible Matter Sources

The eversion of the cell requires a large supply not only of energy, but also of matter. During the membrane transformation process, indeed, the area of the inner membrane must increase by *S*_2_∼ 4*π*^2^*R*^2^ *h*⁄*d*, where *h* ∼ 2.5 *nm* is the distance between the centers of inner and outer layers and *d* ∼ 0.8 *µm* - the minor diameter of the torus (see Fig.1 and Appendix). A further question then arises: what kind of matter source can provide *S*_2_⁄2 *πR*^2^∼ 2 *πh*⁄*d* ∼ 2% of area growth for the inner membrane hemisphere?

The video-microscopy study [8] identified three stages of iRBC opening: (1) pore formation; (2) membrane curling to the maximum pore opening, i.e. to the radius of the cell itself; (3) complete eversion (buckling) of the membrane, which pushes the merozoites into the blood. A quantitative model of the stage 3 is described in detail by Callan-Jones et al. [4]. Our equations describe the energy conversion mechanism of the stage 2 (cell curling) driving the flow of energy and matter from the overstressed inner layer. This curling is the key in the egress process because during this stage the iRBC is irreversibly destroyed and releases the merozoites. The dynamics of the stage 3 should also fall into the physical scenario under consideration, but equation (5) for pore growth is no longer applicable when the pore size has stabilized.

To realize a scenario of the cell eversion, described above, the inner membrane layer of the infected cell needs to accumulate energy and matter to prepare egress as an instability and an irreversible phenomenon triggered by the pore opening. In principle, the accumulation of energy in the internal leaflet can be due to its saturation by additional molecules [27] or by interaction of the membrane with proteins. Merozoites, penetrating into the erythrocyte, then develop inside the parasitophorous vacuolar membrane (PVM). Before eversion of iRBC and egress of merozoites, PVM is destroyed [1] so as not to interfere with the release of parasites. It is not yet clear whether PVM disruption plays any role in the cell opening process. Possibly, PVM residues can be used by merozoites to additionally saturate the inner layer of the iRBC membrane with lipids. The present level of knowledge is not sufficient to rule out any of these mechanisms and a comparative analysis of all possible mechanisms is beyond the scope of this article. Here, we provide and explore a more detailed quantitative evaluation of the mechanism based on a hypothesis that matter is brought to the inner membrane of iRBC by macromolecules and lipid vesicles produced by the parasite (Figure 3). This produces an excess area of the internal leaflet, accumulating an internal mechanical stress ready to be released as soon as the pore opens. Vesicles in Figure 3 can mean single membrane structures like micelles or Maurer’s cleft, as well as bilayer vesicles, if there is a mechanism for their preferential attachment to the inner layer of the cell membrane. Membrane fusion depends on many factors [28], but the complexity of the fusion process is beyond the scope of the manuscript.

**Figure 3.**
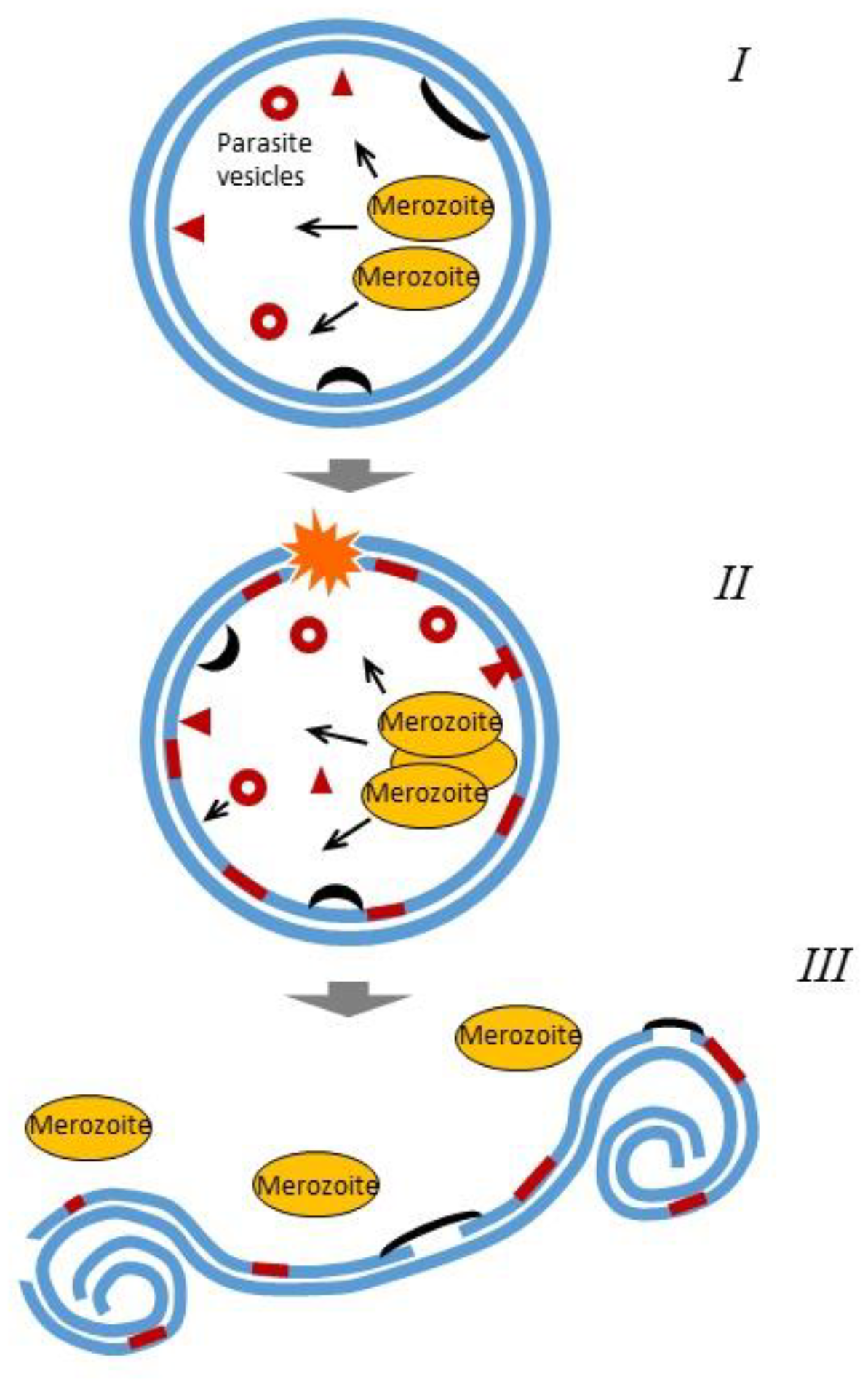
Schematic representation of different stages of iRBC eversion and egress of malaria merozoites within the suggested mechanism. From top to bottom: (i) generation of vesicles (circles) and molecules (arrows) by merozoites (ii) fusion of the merozoites with the inner leaflet of iRBC membrane, which increases the excess of lipids in the inner leaflet. Pore formation and (iii) curling and final eversion of iRBC. We assume that remnants of the cytoskeleton (black arches) may be involved in supplying the inner membrane with energy, which can lead to the membrane rupture.

### 3.5. iRBC eversion scenarios

From this perspective, we can consider two limiting scenarios for iRBC eversion.

Scenario A: with maximum energy consumption (*E*_*max*_ ∼10^−15^ *J* per erythrocyte), which is used both to change the shape and curvature of the membrane, and to tear the inner layer of the membrane.

Scenario B: with minimal energy consumption (*E*_*min*_ ∼ (1 − 2) ∗ 10^−16^ *J* per erythrocyte), which assumes that no ruptures of the inner layer of the membrane occur. This scenario requires an additional source of lipids to be incorporated into the inner membrane of the cell as it stretches.

According to scenario A, let us estimate how many molecules need to be attached to the inner layer of the membrane in order to obtain the energy *E*_*max*_ ∼10^−15^ *J* required for the eversion of iRBC:

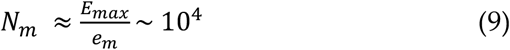

where *e*_*m*_∼10^−19^ *J* – the elastic energy per lipid molecule [28]. According to Helfrich (1973) [28], indeed, one can estimate *n*∼10^19^ *m*^−2^ – as the number of molecules per square meter of a membrane and *n* ∗ 4*πR*^2^∼10^9^ – the total amount of molecules in one leaf of the membrane. Consequently, 10^4^ molecules add only 10^−3^% of the surface area of the membrane, which is noticeably less than 2%, which is needed to transform the membrane without rupture.

The scenario B of cell transformation with minimal energy consumption is possible if we consider the option of attaching not individual molecules to the inner cell membrane, but macromolecular vesicles. Such vesicles carry not only elastic energy, but also a significant amount of lipids, which allows the inner leaf of the membrane to expand without rupture. Let us estimate how many vesicles are needed to deliver the required amount of matter and energy to the inner layer of the membrane:

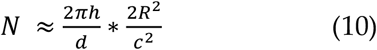

where 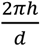 is about 2% of excess area of the inner membrane leaf, *R* ∼3 *µm* is the red blood cell radius, *c* is the diameter of the vesicle. These vesicles contain an amount of elastic energy equal to 2*π*κ*N* (see Appendix). Thus, if the energy stored in vesicles will be released for the diffusion time *t*_*d*_, it provides support by energy for the cell transformation at the scale:

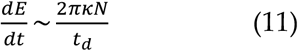

If we replace the value of *t*_*d*_ with the time of curling *t*_*curl*_ in equation (11), we will get:

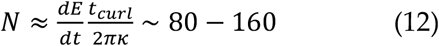

The typical size of these vesicles can be estimated from Eq. (10). One iRBC gets sufficient number of lipids for eversion from *N*∼80 − 160 vesicles with typical diameter 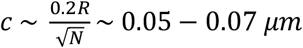. An intermediary scenario between A and B can be imagined, when energy is delivered to the inner membrane mainly by molecules, and matter is delivered by a few large vesicles.

### 3.6. Experimental evidence

The participation of vesicles in the eversion of iRBC is confirmed by experiments. Microscopic studies show clusters of vesicles localized near the iRBC membrane as well as the characteristic concavity or alveoli of the cells, probably associated with the attachment of the parasite vesicles to the membrane [30, 31]. The parasites *P. ovale* can use the variant of vesicles known as Schüffner’s dots [30, 31]. Other types of parasites can choose the variant of saturation of the membrane with individual molecules.

In the case of *P. ovale*, from the microscope images we can estimate the number of Schüffner’s dots in each iRBC as ∼100 with a diameter of ∼0.1 µm [32]. In the case of *P. falciparum*, uncoated vesicles with a diameter of 0.025 *µm* and coated electron-dense vesicles with a diameter of 0.08 *µm* have been described [33, 34]. The 0.025 *µm* vesicles transfer matter (lipids and/or proteins) from the Maurer’s clefts to the inner membrane of the iRBC [31, 34-36]. Although we consider a spherical shape for the vesicles, a slight change of the shape does not significantly affect the results. Thus, the quantitative estimation presented here shows that our theoretical model for the vesicles-mediated mechanisms of iRBC eversion is in good agreement with experimental data.

These considerations suggest the following potential scenario of the iRBC opening (Figure 3): 1) Parasites get into the RBCs and create specific organelles (Maurer’s clefts, Schüffner’s dots and electron-dense vesicles); 2) These organelles produce an excess of matter in the inner leaflet of the iRBC membrane and, consequently, its mechanically stressed state with respect to the external leaflet. This can be done by addition to the inner leaflet of either individual lipids [37] or knobs [9]. The excess of energy and matter is irreversibly released after the pore opening in the iRBC. 3) The cell with the inner leaflet with an excess of matter, thus under mechanical internal stress, store mechanical energy ready to be released with an energetically favorable process via curling and eversion of the cellular membrane. This scenario is supported by numerous microscopic studies revealing several types of vesicles and clefts generated by the parasite. Among them are Maurer’s dots and clefts, Schüffner’s dots or vesicles generated after the distraction of parasitophorous vacuole membranes [9, 33-40]. Note that our model suggests a possible function of Maurer’s dots and clefts, Schüffner’s dots and various vesicles, which remains yet a mystery in parasitology [36].

We are aware that the detailed molecular mechanism of the vesicle fusion needs further investigation. The fusion of the lipid layers generated by the parasite with the iRBC membrane most probably requires the participation of proteins for lipid insertion against an excess of lipids in the inner membrane. Indeed, a number of studies described several proteins that may affect the process of the iRBC eversion [41-46]. Antibodies to PfSEA-1 decreased parasite replication by arresting schizont rupture [41]. Parasites lacking SERA5 undergo accelerated but defective egress in which the red cell membranes do not rupture properly [42]. The paper [46] considers the possibility of the chemical modification of an existing active-site inhibitor of falcipain, which can block hemoglobin degradation by the malaria parasite. Our model considers the mechanical and hydrodynamic aspects of cell eversion. A discussion of the detailed chemical impact on these processes is beyond the scope of our article, but we believe that building a correct physical model for the release of merozoites from an infected cell will help find effective chemical methods for influencing the process of merozoite reproduction. For example, with a decrease in the energy reserve on the inner layer of the cell membrane, the number of unopened cells should increase significantly (experimental data usually indicate few percent of unopened iRBCs [8]).

## 4. Conclusion

Models describing an infected erythrocyte usually assume that the excess elastic energy stored in the compressed inner leaflet of the vesicle is actually driving the curling process. In previous studies [4, 8] on the eversion of infected erythrocytes, it has been considered that the inhibitory factor determining the rate of eversion is viscosity. In this paper, we show that a more important inhibitory factor can be the formation of a rim at the edge of the pore, while the rate of the membrane curling after the rim opening is controlled by the lateral diffusion of lipids.

We described a quantitative model of iRBC eversion during egress of malaria parasites. From our estimate, we found that this transformation of the cell requires an increase in the area of the inner cell membrane of about 2% and the minimal energy of *E* ∼ 10^−16^ *J*. The model leads to a simple formula explaining the experimentally observed speed of the iRBC curling. We thus show that medium viscosity does not play a significant role in iRBC eversion. We hypothesize that this process is actually controlled by the lateral diffusion of lipids.

Our level of knowledge is not sufficient to unambiguously state that the saturation of the inner membrane of iRBC, by an excess of energy and matter, is due to additional lipid or (and) protein molecules. Here, we evaluate in detail one of the mechanisms, which is based on the involvement of vesicles of the parasites. Our quantitative estimations suggest that this mechanism is possible, although scenarios involving molecules or cytoskeletal residues are also physically consistent. This mechanism may shed light on the role of the parasite-generated structures like Maurer’s cleft and Schüffner’ dots, whose exact functions are still unknown. Finally, the vesicle-mediated mechanism can be extended to other cellular processes [45] where compact vesicles can serve as a source of energy and matter to drive further mechanical instabilities.

This quantitative model of curling of the iRBC membrane during egress of malaria parasites also provides a framework for further tests and comparative analysis with other possible mechanisms as soon as new experimental data will become available. Indeed, based on the equation (11) describing the speed of iRBC transformation, we can predict that the rate of iRBC transformation and the release of merozoites could be slowed down or even stopped by different mechanisms. To cite few examples: (i) the increase in the lateral diffusion coefficient of lipids (i.e. by increasing the cholesterol concentration and by lowering the temperature); (ii) the reduction of the number of vesicles and clefts inside iRBC (or even the inhibition of their production); (iii) the inhibition of the attachment of vesicles and clefts to the inner lipid layer of iRBC and (iv) the inhibition of the machinery for lipid insertion in the inner membrane leaflet. These predictions can be tested experimentally and may lead to new approaches to prevent malaria.

## Author Contributions

NG developed the theoretical formalism and wrote the initial version of the manuscript, AP discussed the model hypothesis and verified the results, AVK supervised the project. NG, AP and AVK discussed the results and contributed to the final manuscript.

## Acknowledgements

This work was supported by the grant of the Region Languedoc-Roussillon for invited professors and was partially accomplished during a monthly stay of N.G. at the University of Montpellier as a Visiting Professor. A.P. acknowledges the LabEx NUMEV (ANR-10-LABX-0020) within the I-SITE MUSE of the University of Montpellier (No. AAP-2016-2-025) for financial support and the CNRS for a dispense of a semester (demi-delegation) from teaching duties. The authors thank C. Braun-Breton, J. F. Dubremetz, M.M. Carretero, A. Vasilkov and N.-O. Walliser for helpful discussions and S. Andreev for critical reading of the manuscript and suggestions.

## Notes

### Competing Interest Statement

The authors have declared no competing interest.

