## Supplemental Data for "Quantitative model and physical mechanisms of iRBC membrane curling during egress of malaria parasites"

### AI. Changing the area of the membrane during cell eversion.

This article is devoted to the process of erythrocyte curling, which takes from the formation of a pore to its maximum opening. Consequently, all equations and estimates are given not for the full sphere, but for half of the sphere (Fig. 4).

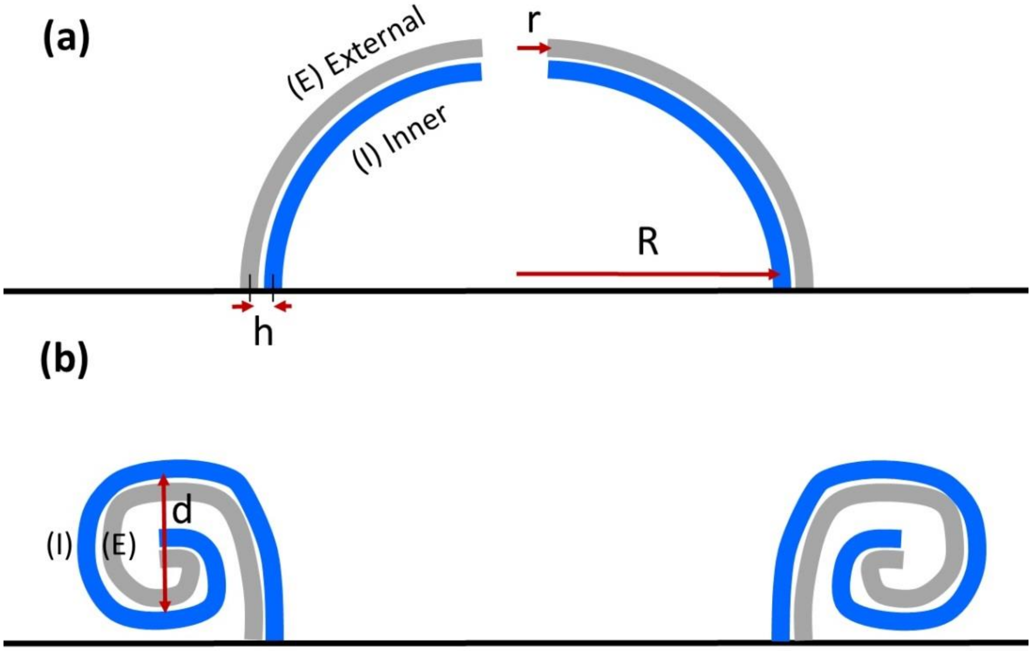


**Figure 4.** Rearrangement of the upper hemisphere of the iRBC (a) when the membrane is inverted. The outer and inner layers of the membrane in the initial state (4a) are marked with the letters (E) and (I).

In the initial state of the cells (Fig. 4a) outer layer area (leaflet) is larger than the area of the inner layer by an amount (for one hemisphere):

$$S_0 \sim 2\pi(R+h)^2 - 2\pi R^2 \sim 4\pi R h \qquad \text{(A1)}$$

During curling of iRBC, the inner leaf of the cell transforms in the external leaflet of the rim. The area of the external layer of a cylinder is larger than the area of the inner layer (Fig. 4b) by the amount of

$$S_1 \sim 2\pi R * \left[2\pi * \left(\frac{d}{2} + h\right) - \pi d\right] \sim 4\pi^2 Rh \qquad \text{(A2)}$$

Equation (A2) corresponds to the case of maximum cell opening when the pore size is r=R. The number of cylinders is $\sim R/d$ (as ratio of perimeters of cross sections), so total increasing of surface of former inner leaf of the cell is

$$S_2 \sim \frac{4\pi^2 R^2 h}{d} \qquad \text{(A3)}$$

where $R \sim 3\ \mu m$ is the radius of the cell, $h \sim 2.5\ nm$ is the distance between the centers of lipids from inner and outer layers, $d \sim 0.8\ \mu m$ - the minor diameter of the torus. From (A1) and (A3) one gets $S_2 \gg S_0$ , because $\frac{\pi R}{d} \gg 1$. The growth of area of the inner membrane (leaf) is $S_2/2\,\pi R^2 \sim \frac{2\pi h}{d}$ or ~2 % of the initial surface of a cell (for one hemisphere).

This area increase of the inner membrane is only possible in two cases: 1) the inner membrane had an additional source of matter, such as lipids, which is used for its expansion; 2) the inner membrane was ruptured when the erythrocyte was everted. The formation of such gaps requires a significant expenditure of energy, which must be stored in the inner layer of the cell membrane. Note that the energy spent on breaks can exceed the energy required to transfer the cell from state 4a to state 4b, which has a large elastic energy (assuming the same local properties of the membrane). It is also obvious that it is impossible to rearrange the erythrocyte membrane from state 4a to state 4b, when the initial pore expands to the cell diameter, without lateral diffusion of lipids along the inner and outer layers of the membrane. This means that the characteristic time of lipid diffusion can play an important role in the dynamics of erythrocyte membrane rearrangements.

The time between membrane lipids exchange layers exceeds six hours (see, for example, review by Van Meer, 2011 [25]) and even longer (days). Thus, exchange of substances between the layers during rapid cell curling can be ignored, although partial leakage of lipids from the inner layer to the outside layer could occur during the 48-hour cycle of the parasite, probably partly responsible for the conversion of normal disc-shaped in a spherical infected erythrocyte.

**AII. Elastic energy**

A cell membrane with spherical shape has a specific (per unit area) elastic energy (see, for example, Abkarian at al, 2011 [8]):

$$E_m = \frac{\kappa}{2a^2} \qquad \text{(A4)}$$

where $a$ - the radius of curvature of the membrane. The more curved membrane and the smaller the radius $a$, and the more it comprises elastic energy per unit area. Most elastic energy stored in the rim. By neglecting the major curvature radius of the toroidal rim (i.e. we approximate its energy by that one of a cylinder), the elastic energy can be written as:

$$E_i \approx \frac{\kappa}{2{d_i}^2} * \pi d_i * 2\pi r = \frac{\pi^2 \kappa r}{d_i} \qquad \text{(A5)}$$

For $d_i = 2iL; i = 1, \dots, N_c$ , where $N_c$ – the number of the cylinders in the rim, the sum of energy of all cylinders:

$$E_{rim} \approx \sum E_i \approx \frac{\pi^2 \kappa r}{2L} \sum 1/i \qquad \text{(A6)}$$

For $N_c \approx 4$ (from experimental data by Abkarian at al, 2011) we can estimate $\sum 1/i = 1 + \frac{1}{2} + \frac{1}{3} + \frac{1}{4} + \cdots \approx 2$ . The elastic energy stored in the tightly folded multilayer membrane can be estimated

$$E_{rim} \sim \frac{\pi^2 \kappa}{L} r \qquad \text{(A7)}$$

Total elastic energy of a spherical cell (or vesicle) with radius $R$ can be calculated from the formula of the specific energy and the cells surfaces:

$$E_{sph} = \frac{\kappa}{2R^2} 4\pi R^2 = 2\pi\kappa \qquad \text{(A8)}$$

Comparing (A7) and (A8), we find that for $r \gg 0.1\ \mu m, \pi r/(2L) \gg 1$. Consequently, $E_{rim} \gg E_{sph}$.
